# Relevance of network topology for the dynamics of biological neuronal networks

**DOI:** 10.1101/2021.02.19.431963

**Authors:** Simachew Abebe Mengiste, Ad Aertsen, Arvind Kumar

## Abstract

Complex random networks provide a powerful mathematical framework to study high-dimensional physical and biological systems. Several features of network structure (e.g. degree correlation, average path length, clustering coefficient) are correlated with descriptors of network dynamics and function. However, it is not clear which features of network structure relate to the dynamics of biological neuronal networks (BNNs), characterized by non-linear nodes with high in- and out degrees, but being weakly connected and communicating in an event-driven manner, i.e. only when neurons spike. To better understand the structure-dynamics relationship in BNNs, we analysed the structure and dynamics of > 9, 000 BNNs with different sizes and topologies. In addition, we also studied the effect of network degeneration on neuronal network structure and dynamics. Surprisingly, we found that the topological class (random, small-world, scale-free) was not an indicator of the BNNs activity state as quantified by the firing rate, network synchrony and spiking regularity. In fact, we show that different network topologies could result in similar activity dynamics. Furthermore, in most cases, the network activity changes did not depend on the rules according to which neurons or synapses were pruned from the networks. The analysis of dynamics and structure of the networks we studied revealed that the effective synaptic weight (*ESW*) was the most crucial feature in predicting the statistics of spiking activity in BNNs. *ESW* also explained why different synapse and neuron pruning strategies resulted in almost identical effects on the network dynamics. Thus, our findings provide new insights into the structure-dynamics relationships in BNNs. Moreover, we argue that network topology and rules by which BNNs degenerate are irrelevant for BNN activity dynamics. Beyond neuroscience, our results suggest that in large networks with non-linear nodes, the effective interaction strength among the nodes, instead of the topological network class, may be a better predictor of the network dynamics and information flow.

## Introduction

Complex networks have become an essential mathematical framework to understand the dynamics and information flow in many natural and social systems. One of the key questions in network science is to understand how network structure shapes network dynamics. Several measures of network connectivity structure have been identified to describe the dynamics and information flow in a network. Moreover, it is argued that certain mathematically well-defined complex random networks (e.g. Erdös-Rényii (ER), small world (SW), scale free (SF)) have different dynamics, robustness and information flow properties [1].

Different network topologies may render the network with different properties. For example, ER networks are usually more robust to node and edge degenerations and generalize well, due to the homogeneity of connectivity of nodes [2]. On the other hand, SW topology facilitates better segregation and integration of information in the network, due to its higher clustering and shorter path length property[3, 4]. The hub nodes in scale-free networks are more important in spreading information rapidly and in keeping the network intact upon neurodegenerative assault [4, 5].

In a real-world scenario, both natural and social systems are highly dynamic. That is, both the number of interacting nodes and their connections change over time. Neuronal networks in the brain are excellent examples of such dynamic networks, where the topology, size and connection density are continuously modified by normal (e.g. development, synaptic plasticity, learning) or abnormal (e.g. neurodegenerative diseases) processes.

The topology of the brain networks depends on the spatial scale. Within a brain region at short distances (<100 *µ*m), the connectivity appears to be random (ER). At mid-range distances (0.1 − 10 mm), the connectivity is random, but the connection probability decreases with distance [6, 7]. Finally, at a large scale (>1 cm), connections often appear to follow the scale-free or small-world topology [4]. In neurodegenerative diseases such as Parkinson’s disease and Alzheimer’s disease, both neurons and/or synapses die and the networks lose their functions. To understand the functional deficits associated with such neurodegenerative diseases, it is of key importance to understand how the loss of neurons and synapses affect the network activity dynamics.

Therefore, here we address two key questions: 1. How are the activity dynamics of biological neuronal networks (BNNs) altered when either neurons or synapses are removed from the network? 2. Can we see signatures of the network topology in the network activity dynamics? To address these two questions we analysed the dynamics of BNNs with three well-studied network topologies: random [8], small-world [9], and scale-free [10]. We perturbed each of the three network topologies by pruning either the edges (synapses) or the nodes (neurons). Surprisingly, we found that the network topology was not a good predictor of how the network activity evolved as neurons and synapses progressively died. Moreover, the rules that governed the death of neurons or synapses did not have any qualitative effect on the network activity dynamics. Further analyses of network structure and network dynamics of 9,090 different networks revealed that for BNNs, the effective synaptic weight (another measure of network structure) was a better predictor of the network activity dynamics than the network topology.

## Results

### Network activity states do not depend on network topology

To study the effect of network topology on the activity dynamics, we considered directed networks with common topologies; namely, ER, SW and SF (cf. Methods, Fig. 1). To expose the differences among these network topologies, we systematically deleted either edges (synapses) or nodes (neurons). In each network, the nodes were modelled as leaky-integrate-and-fire neurons of which 80% were excitatory and 20% inhibitory, to mimic the fractions of these neurons in the neocortex (cf. Methods). We varied the ratio of total inhibitory and excitatory input (*g*) to the neurons to systematically change the excitation-inhibition (E-I) balance and to tune the network in different activity regimes [11, 12]. In addition, all neurons received Poisson-type spike trains as external excitatory input. To create a perturbation in the network structure, we designed five different neurodegenerative and five synaptic pruning strategies [13]. Finally, to quantify the network activity states for all these different networks, we estimated both single neuron activity descriptors (mean and variability of firing rate, spike time irregularity) and population activity descriptors (average pairwise firing correlation and population firing asynchrony) (cf. Methods; see Fig.2 [11, 12]).

**Fig. 1.**
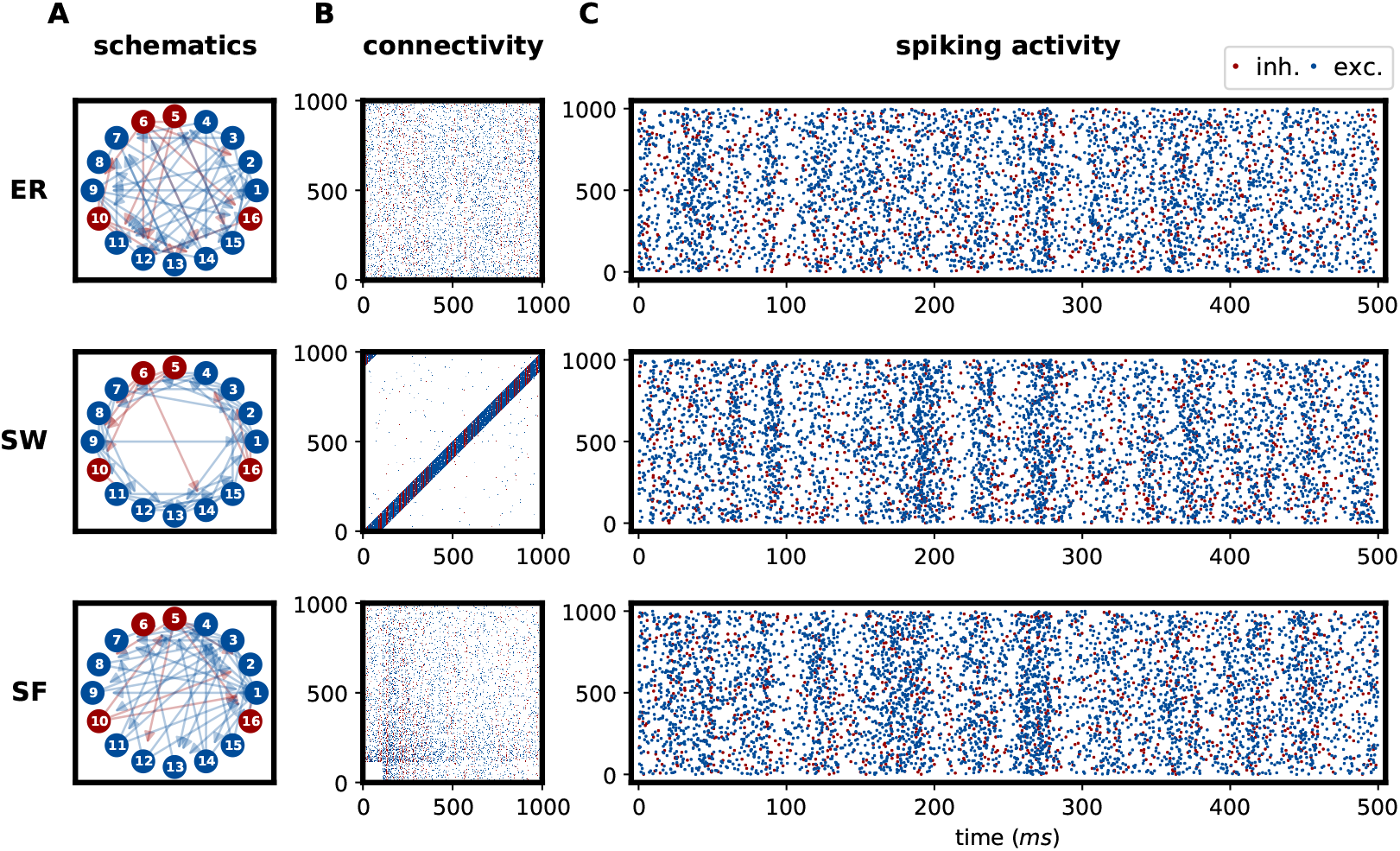
Network connectivity and spiking activity of parent networks. A. Schematic of three types of network topologies. B. Example of adjacency matrices of the three types of networks. Here we used 1,000 neurons (800 excitatory and 200 inhibitory) which were sparsely connected (with 10% connection density in their unpruned state). C. Spike activity raster for the networks shown in panel B. Here the networks were tuned to operate in an inhibition dominated regime *g* = 5.

A comparison of the network activity states (an example is shown in Fig.1) revealed that all three network types had similar neuronal activity statistics in different activity regimes, regardless of the differences in their network topologies. This similarity was surprising, but it could be a consequence of our particular choice of inputs, connection probabilities and E-I balance., We hypothesized that differences in these network topologies could be revealed if we perturbed the network connectivity by pruning either the synapses or the neurons, as each network topology should evolve along its specific trajectory as we systematically pruned the network. To test this hypothesis, we degenerated the network using five synaptic pruning [13] and five neuronal pruning strategies (cf. Methods). Fig. 1 shows how the five synaptic pruning strategies changed the structural and functional features of an ER network for *g* = 5 to illustrate the steps of our analysis.

In both inhibition-dominated regimes (*g* = 5 and *g* = 7, Fig.2, top and bottom rows), the firing rate increased (Fig.2A, bottom row), spike time irregularity decreased (Fig.2B, middle and bottom rows) and synchrony decreased (Fig.2C, middle and bottom rows) as the networks degenerated, irrespective of the network topology and pruning strategy, either by synaptic pruning (left-half of each panel in Fig.2) or by neuronal death (right-half of each panel in Fig.2). In these inhibition-dominated regimes, recurrent inhibition is responsible for keeping the network activity low, irregular and uncorrelated. Because the network degeneration globally reduced the level of inhibition in the inhibition-dominated regime, the firing rate increased with synaptic loss and neurodegeneration. In this regime, all networks showed only weak correlation, even in their unpruned state. Hence, there was only a small change in the network synchrony as we degenerated the networks, converging onto the completely uncorrelated state driven by uncorrelated Poisson inputs. Thus, surprisingly, neither network topology nor synapse pruning strategy made any difference to degeneration-induced changes in the network activity state. Instead, all pruning-induced changes could be explained by how pruning altered the E-I balance and sparsity of the network. Similar results were also observed for the excitation-dominated regime of the network activity dynamics (see Supplementary Fig. S1). Because the excitation-dominating regimes have little biological significance, we focused on the inhibition-dominated regime.

**Fig. 2.**
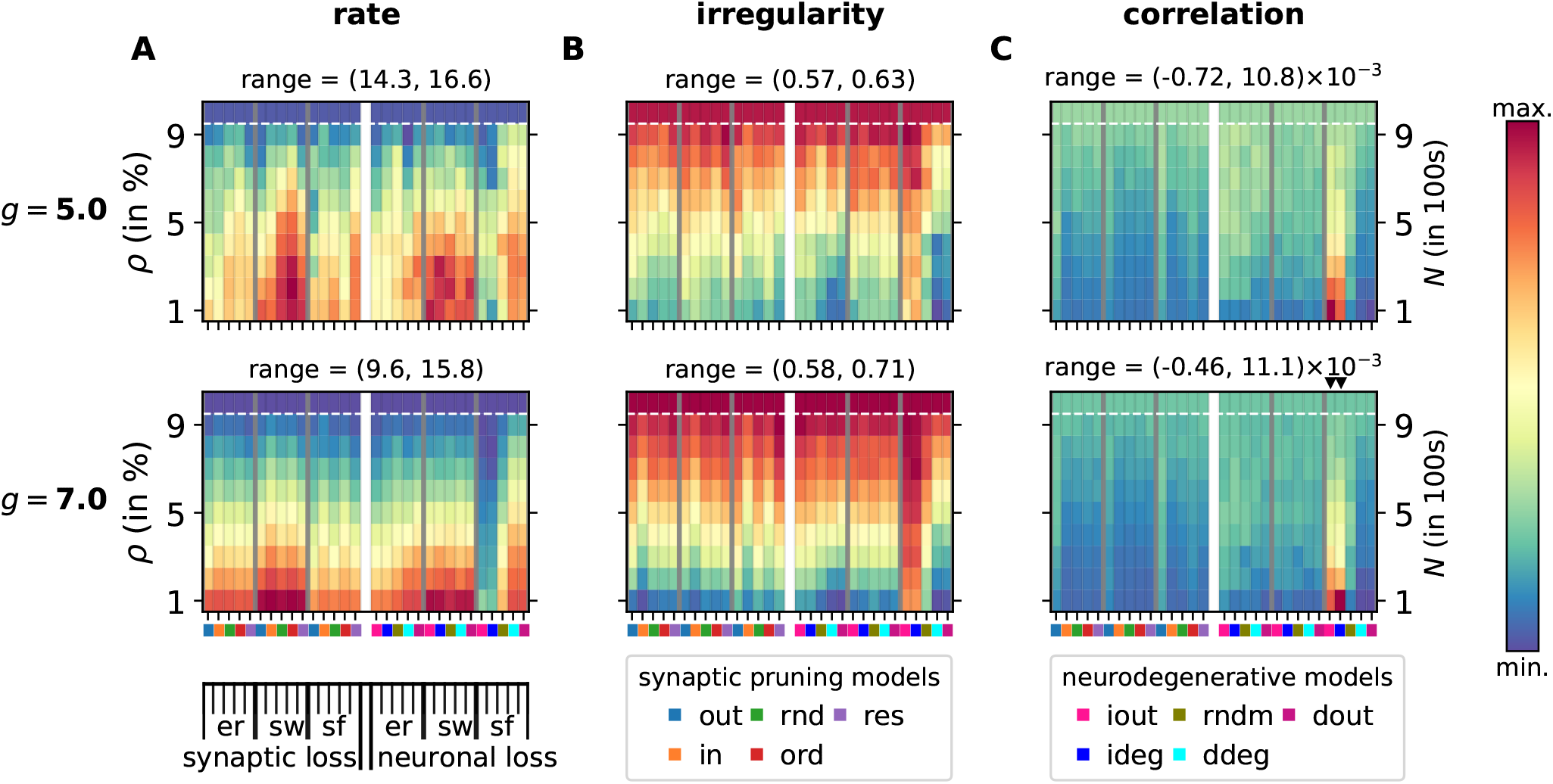
The changes in the activity states of neuronal networks due to different types of degeneration. Each panel is divided by a white vertical midline to separate synaptic pruning on the left half and neurodegeneration on the right half. The vertical axis for the left half denotes connection density *ρ*, for the right half it denotes network size, *N*. The degeneration progresses from top to bottom. Each half is further subdivided into three partitions, corresponding to ER, SW and SF networks. Each partition comprises the five synaptic pruning strategies on the left half and the five neurodegenerative strategies on the right half. The synaptic pruning strategies are arranged as ordered, out-, random, resilient and in-pruning strategies, whereas the neurodegenerative strategies are arranged as increasing-out-degree, increasing-degree, random, decreasing-degree and decreasing-out-degree degeneration strategies (see the color code for these at the bottom of the Figure). Each small square in the colormaps represent an average value (10 realizations) of the specified network activity descriptor. Top and bottom rows represent weak (*g* = 5) and strong (*g* = 7) inhibition-dominant states. Mean firing rate (**A**), spike time irregularity (**B**) and spike time asynchrony (**C**) of the networks undergoing progressive synaptic pruning and neurodegeneration for the two activity regimes.

Although pairwise correlations generally decreased with degeneration, we observed two exceptions. For the scale-free networks, pairwise correlations increased for two of the neurodegenerative strategies (iout and ideg; cf. Methods) as we reduced the number of neurons in the network (Fig.2C - right half, columns marked by downward triangles). In these two neurodegenerative strategies, we primarily removed neurons that were weakly connected. This preserved the hubs at the later stage of degeneration, thereby facilitating high shared connections leading to increased pairwise correlations. These results suggest that, barring these two exceptions, the network activity state was neither influenced by network topology, nor by the specific neuron or synapse pruning strategy.

### Effect of degeneration on network topology

To understand why network activity was not influenced by the different network topologies and degeneration strategies, we quantified how our different degeneration strategies altered the network topology. To this end, we estimated the connection density, the number of pairs that shared common presynaptic neighbours, spectral radius, node centrality, clustering coefficient and characteristic path length as pruning progressed in each of the network topologies (cf. Methods). In addition, we also considered the degree distribution and computed the topology indices of the networks (randomness, small world index, scale-freeness; cf. Methods) to label them as random, small-world, scale-free or any other type.

By design, all synaptic pruning strategies removed equal numbers of synapses at each stage of pruning. Therefore, the connection density and node degree were preserved for all three topologies at each pruning stage. However, other measures of network structure changed in a similar fashion for all three network topologies (Fig. 3 A). Progressive synaptic pruning and neurodegeneration both clearly withered the node centrality in a monotonic fashion for all three network topologies (Fig. 3 A, left). However, the relative decrease in node centrality differed for each neurogenerative strategy. Whereas two of our non-random neurodegenerative strategies (dout and ddeg) preferentially pruned neurons with higher degree, the other two (iout, ideg) removed neurons with lower degree. As expected, networks lost their node centrality faster when they lost neurons with higher degree (Fig. 3A - left).

**Fig. 3.**
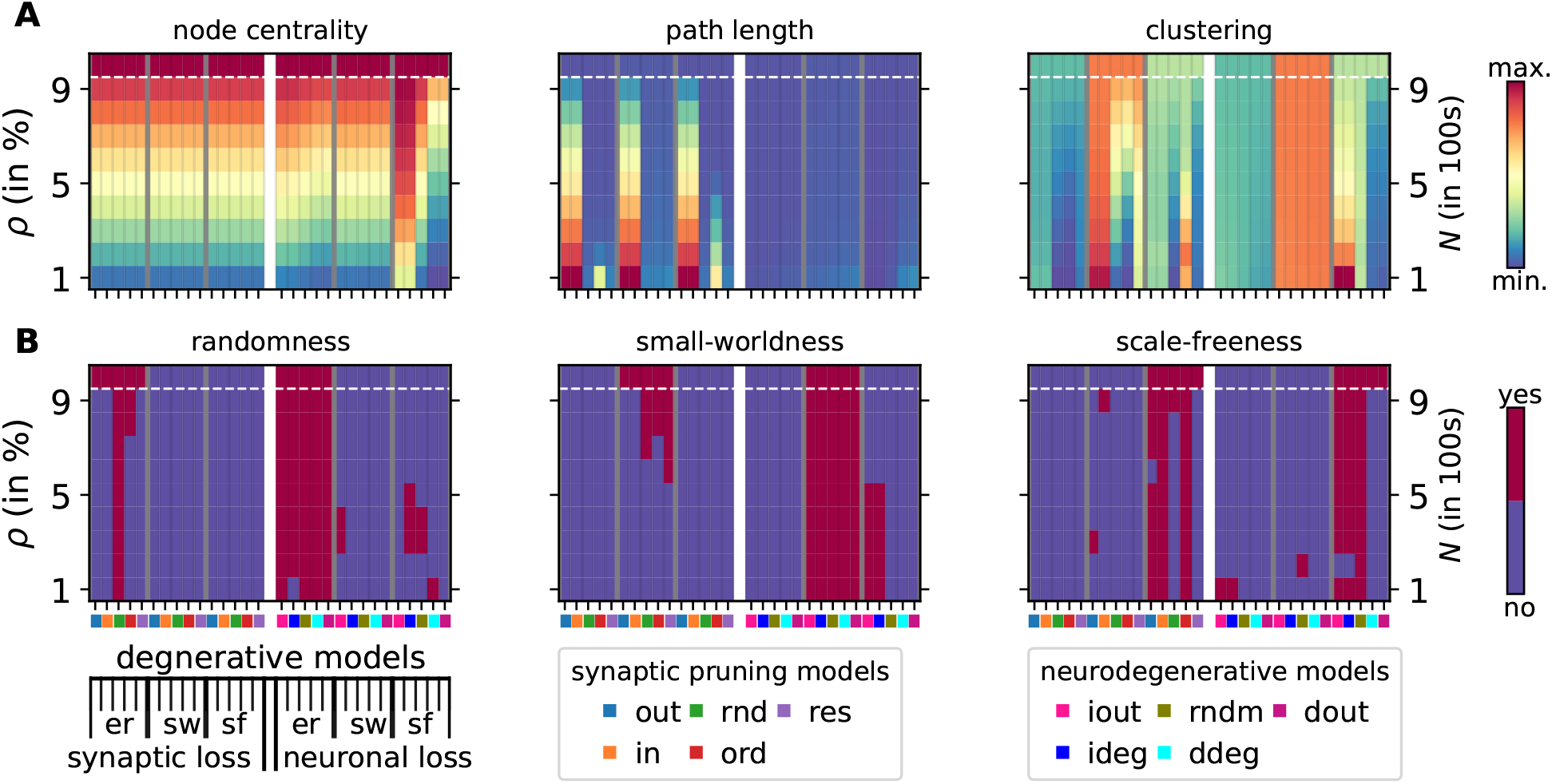
Changes in structural features of neuronal networks due to different types of degeneration. Each panel is presented in the same way as in Fig.2. **A** Different measures of network structure: node centrality (left), average shortest path length *L* (middle), and clustering coefficient *cc* (right) of the directed networks. Normalization was performed across parent networks. **B** Changes in the network topology with progressing degeneration. Red color indicates that the network satisfies the criteria for ER (left), SW (middle) and SF (right). Randomness was estimated using a Chi-squared (*χ*^2^) test for randomness using a Gaussian degree distribution fit. Small-worldness was estimated using the small-world propensity condition (*ϕ* >= 0.6), computed from path-length and clustering. Scale-freeness was defined by the extent of fulfilling a power-law degree distribution (cf. Methods).

Independently of the network topology as well, both synaptic and neuron degeneration resulted in an increase in the average characteristic path length, because of the loss of short-cuts (Fig. 3A - middle). While there were not differences associated with the network topology, two synaptic pruning strategies (out-pruning and in-pruning) increased the average characteristic path length of the network more significantly than the other pruning strategies (Fig. 3A - middle). This was because these two pruning strategies prioritized the sequential removal of all outgoing or incoming projections of the neurons, respectively. In contrast to the synapse pruning, our neurodegenerative strategies did not affect the average shortest path length, because neuron loss also meant synapse loss. Whereas neuron loss by itself decreased the path length, the concomitant synapse loss increased the average path length.

Similar to characteristic path length and node centrality, network degeneration also reduced clustering in the networks, again irrespective of their network topology (Fig. 3A - right). The pruning induced decrease in the clustering coefficient was smaller in SW networks as these networks have higher clustering to begin with. In fact, clustering is the main property that defines a SW network [9]. Note that clustering is not always expected to decrease by synaptic loss, because clustering is an estimate of how well the neighbours are themselves interconnected. In fact, two networks of the same size but with different connection densities can have equal clustering coefficients. For example, if a neuron has only two synaptic neighbours, its clustering coefficient would be maximal if the neighbours were themselves interconnected, whereas a hub in a sparse network would have a small clustering coefficient if the neighbours were not well interconnected.

Among the five neurodegenerative strategies, only the random loss of neurons preserved the clustering in all networks. The two neurodegenerative strategies that progressively targeted poorly connected neurons (iout and ideg) increased the clustering for all network types. The neurodegenerative strategies that removed neurons with relatively higher degrees (dout and ddeg) increased the average clustering in small-world networks. Because, by definition, SW networks have high clustering and homogeneity in degree distribution, neurodegenerations in increasing or decreasing degree had a similar impact on the network. By contrast, in ER and SF networks, neurodegeneration resulted in rapid loss of clustering. Moreover, in SF networks, neurodegeneration also removed hubs.

To further characterize how synapse and neuron loss altered the network structure, we estimated how random, small-world or scale-free the networks were after pruning (cf. Methods). Our analysis revealed that in most cases, synapse or neuron pruning altered the structure of the network topology qualitatively, such that the network failed to maintain their original characteristic topology in the wake of neurodegeneration (Fig. 3B). Given that each network topology was altered following 10 different degeneration strategies, ER networks maintained their randomness for three, SW networks maintained their small-worldness for five and SF networks maintained their scale-freeness for six strategies. SW topology was most robust to neuron loss, whereas SF topology was more robust to both synapse and neuron loss.

### Effective synaptic weight is a better indicator for network dynamics

The aforementioned analysis suggests that network topology by itself is not an important descriptor of the network activity dynamics. That is, network structure descriptors such as clustering coefficient, characteristic path length, etc. are not particularly useful for determining the dynamics of neuronal networks. This raises a pertinent question: which graph theoretic descriptors are more useful for determining the network activity dynamics? In the previous Sections, we generated 5,460 networks with different connection structures, operating in different activity regimes, to characterize the effect of network degeneration on structural features and network activity. With 3,600 additional networks, we identified the features of network structure most correlated with the network activity dynamics.

We used standard multivariate linear regression to identify the contributions of nine different descriptors of network structure to characterize the network activity dynamics (cf. Methods). We found that the effective synaptic weight 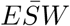 was the most significant measure to infer most network activity statistics (Fig. 4 B, bottom). Key descriptors of network activity dynamics, i.e. average firing rate, variance of firing rates (*σ*_*λ*_), population synchrony, and spike time irregularity revealed a clear positive or negative relationship with 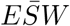 (Fig. 4A, B, bottom). *ESW* essentially estimates the presynaptic connection probability and connection strength. Only the pairwise correlation (*c*) in spiking activity was an exception: it was better predicted by the level of shared presynaptic neighbours and the connection density of the network.

**Fig. 4.**
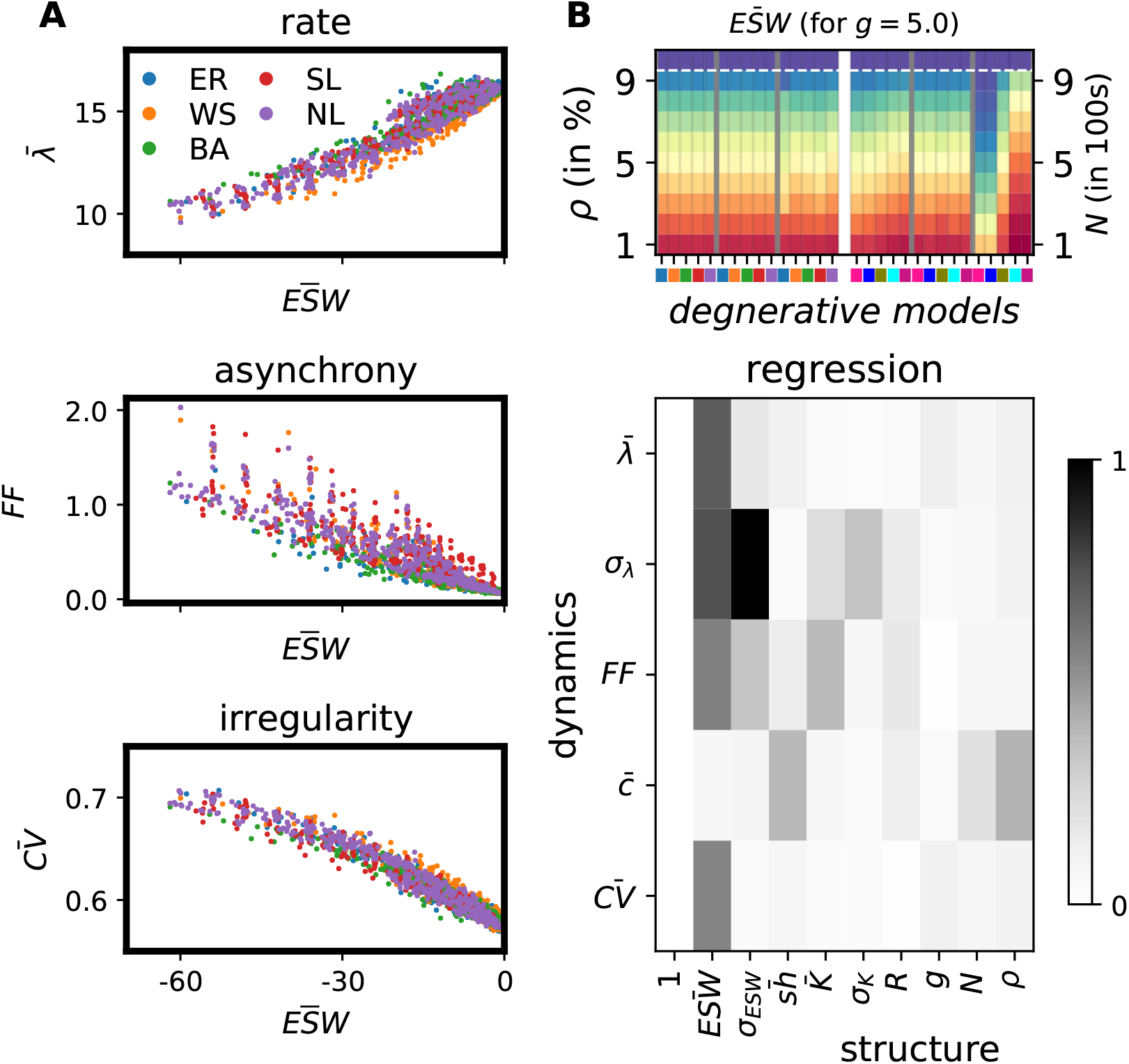
The prominence of effective synaptic weight (*ESW*). A. *ESW* correlated well with the global firing rate (top), asynchrony (middle) and irregularity (bottom). B (top). Changes in *ESW* with degeneration of random, small-world and scale-free networks by five synaptic pruning and five neurodegenerative strategies. B (bottom). Estimation of the five dynamical descriptors from the nine structural quantities of the networks. The dynamical quantities are mean and standard deviation of the firing rates (*λ, σ*_*λ*_), the Fano factor to measure synchrony (*FF*), the mean pairwise correlation of spike trains (*c*), and the coefficient of variation of the inter-spike intervals to measure the irregularity of the spike patterns. The structural quantities are the mean and standard deviation of the effective synaptic weights 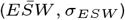, the shared presynaptic neighbours 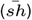, the mean and standard deviation of the degree distribution 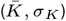, spectral radius (*R*), relative strength of inhibitory synapses (*g*), the network size (*N*) and the connection density (*ρ*). For all structure-dynamics analyses, 300 Erdös-Rényii random, 300 small-world and 300 scale-free networks of 10 different sizes, 10 distinct connection probabilities and 3 different values of inhibition-dominance (*g* = 5, *g* = 6 and *g* = 7) were independently simulated in addition to the degeneration data (cf. Methods).

Finally, we also measured how 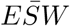 was altered by synapse and neuron pruning. We found that, indeed, 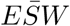 decreased monotonically with pruning, irrespective of network topology and pruning strategy (Fig. 4 B, top). We note here that none of the ten degenerative strategies was devised to systematically alter 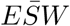 (cf. Methods). Nonetheless, both 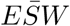 and its variability exhibited strikingly similar changes with network degeneration as major descriptors of network dynamics did, in both excitation-(not shown here) and inhibition-dominant regimes (Supplementary Fig. S2).

## Discussion

Here we studied how the global activity dynamics of BNNs are affected when either synapses or neurons are progressively removed from the network. Specifically, we were interested in understanding whether and how the specific topology of the network and/or the strategy by which it was pruned determined how the network activity changed upon progressive degeneration of its structure. Our analysis revealed two surprising results: 1. The general class of network topology does not influence the evolution of network activity upon network degeneration. 2. The rules according to which neurons or synapses are removed do not influence the evolution of network activity upon network degeneration. According to these findings, the importance of both, network topology and pruning strategy, for predicting network dynamics facing network degeneration is massively overrated. By analysing a larger number of networks with different connectivity structures and activity dynamics we found that for BNNs, the average effective synaptic weight 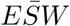 is the most informative structural parameter about the BNN activity dynamics.

To the best of our knowledge, the significance of network topology has not been questioned systematically with one notable exception. Recently, in an attempt to quantify the ’complexity’ of a network, Thikhonov and Bialek [14] also showed that topology is not a significant feature of generic biochemical networks unless it is combined with the connection strength and that interaction strength has a stronger effect than topology on network complexity. Consistent with their results we also argue that for BNNs, effective synaptic strength is a better descriptor of network’s activity dynamics than its topology.

### Topology

Our results do not imply that network topology is a useless feature to infer network dynamics. The three network topologies considered here differed in terms of node in- and out-degree distributions and average path lengths. Hence, the signatures of these topologies should be visible in terms of network synchrony and firing rates. However, in BNNs, neurons have to cross a threshold to elicit a spike and synapses are weak and can be both excitatory (+ve) or inhibitory (-ve). Therefore, neurons need to perform both spatial and temporal integration to elicit spikes and transmit information to their downstream targets. Such spatial and temporal averaging of inputs may reduce the effect of node degree heterogeneity and obscure the network topology differences. In fact, it is known that the out-degree of spiking neurons in BNNs is not a predictor of their impact on the network activity [15].

Another reason why network topology did not make any qualitative difference to the network dynamics is that, as we degenerated the network connectivity, the specific network topology was rapidly destroyed by most pruning strategies (Fig. 3 B). This by itself is not a surprise, because there is usually only one way to construct a network with a given topology and our pruning strategies did not respect that rule. Still, our finding highlights that network topology is not a fixed entity in networks that undergo structural changes, due to functional or dysfunctional reasons.

Instead of network topology, we argue that the effective synaptic weight and its variability (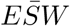 and *σ*_*ESW*_) are better predictors of BNN network dynamics (Fig. 4) and possibly also of other networks that are composed of nodes with a non-linear transfer function and high in- and/or out-degrees.

The interplay of structure and function has long attracted the interest of many studies in neuroscience [15–23]. In particular, there is a great interest in understanding structural correlates of spike synchrony in BNNs. Network structural features such as degree distribution, network motifs, shared inputs and degree correlations have been linked to spike correlations in neuronal networks [21, 24, 25]. Similarly, small-word features play a role in determining local versus global synchronization patterns [26]. Experimental measurements of degree distributions [27], network motifs [28, 29], shared inputs [30, 31] and degree correlations, however, is very tedious. Therefore, we urgently need a better and easier to measure structure-based indicator of network activity dynamics.

Our proposed measure, the effective synaptic weight (*ESW*) provides a better network-structure based indicator for the global network activity dynamics (Supplementary Fig. S2). At the neuronal level, *ESW* measures the capacity of a neuron in receiving and sending information by taking account of the strength and the number of its up- and downstream connections with other neurons (Eq. 5, 6). The variability in *ESW* (Eq. 6) provides a good estimate of how individual neurons differ in their firing activities (Supplementary Fig. S2). For BNNs, *ESW* can be estimated from the intracellular membrane potential of neurons *in vivo*. That is, in a way it is easier to measure *ESW* experimentally, unlike other network topology descriptors which require measurement of the full connectivity of nodes or estimation of the numbers of cycles involving more than two neurons.

### Limitations and future extensions

To reach these results we made several assumptions. Hence, it is important to consider whether our results may change when those assumptions are relaxed. For instance, we only considered networks with rather small connection probability (10%). This choice, however, is consistent with the connection probability measured in the neocortex [19, 32]. As the main emphasis of our work is on BNNs, we are confident that it is not necessary to consider networks with higher connection probabilities. However, to test whether our findings go beyond BNNs, it may be useful to study more densely connected networks in the future.

Next, we only considered networks of moderate size (1,000 neurons). It is non-trivial to extend our result to larger networks (more than 10K neurons), because as we scale the network while keeping the connection probability fixed, we also need to scale the synaptic strengths. That is, for a larger network, the connection strengths may become arbitrarily low and in such a setting it is difficult to predict whether *ESW* will also emerge as the main predictor of network activity. Therefore, in future research it would be interesting to study whether our results also hold for larger networks with appropriate synaptic scaling.

We also assumed that the neurons are inter-connected, independently of their spatial distance. In BNNs with distance-dependent connectivity, the spatial structure of the connectivity can have a big influence on the network activity dynamics [33–35]. Based on our results, we cannot predict how pruning may affect the network ESW when neurons are connected according to their pairwise distances. Hence, this issue requires a detailed investigation on its own.

Finally, we also assumed that the neurons in the network are homogeneous. We chose to study networks with LIF neurons with identical properties, because we wanted to specifically study the effect of network topology. Moreover, previous results have shown that the effect of neuron types depends on the dynamical state of the network [36]. Here, we have investigated the effect of pruning when the networks were operating in an asynchronous-irregular state. In that state, neuron spiking patterns do not seem to affect the network dynamics [36]. It is however, an important question to ask in future studies, how pruning may impact the dynamics of a network when it is operating in another state, e.g. close to the boundary between asynchronous and synchronous states.

### Form, function and degeneration

Besides addressing the fundamental problem of the relationship between network structure and network activity dynamics, we also provided new insights into the mechanisms of neurodegenerative diseases. Brain diseases such as Parkinson’s disease (PD), Alzheimer’s disease (AD) and Epilepsy are characterized by a progressive loss of neurons and synapses. In most cases, the progressive loss of neurons and synapses is both a cause and a consequence of the neurodegenerative disease. For example, functional connectivity analysis suggested that in AD the brain loses its small-world features and signal propagation takes longer than in the normal brain because of longer path lengths [37]. In the last few years, graph theoretical measures of functional and structural connectivity have been suggested to unravel the mechanisms underlying brain diseases [38, 39] and computational models have been used to expose the associated damages in network functions [40, 41]. Our findings suggest that, while obviously neuron and synapse degeneration will affect the network topology (Fig. 3), these topological changes do not have any consequences for the global network dynamics, unless these changes are associated with changes in *ESW* (Fig. 4 and Supplementary Fig. S2). More importantly even, our findings suggest that the specific manner in which neurons or synapses are removed from the network is not relevant for the survival of network dynamics (Fig. 2 and 3). Thus, we propose that, instead of estimating network topology changes due to degeneration, the measurement of *ESW* would be a more effective biomarker for diagnosing neurodegenerative diseases.

## Materials and Methods

### Networks

We used three different network topologies to study the effect of network degeneration (synaptic pruning and neurodegeneration) on the dynamics of the network activity: Erdös-Rényii (ER), small-world (SW) and scale-free (SF) networks. We generated the ER and SW networks following the Watts-Strogatz algorithm [9]. ER and SW networks corresponded to 100% and 2% rewiring probability, respectively. Because we required directed networks, the rewiring was performed for both sources and targets of each edge. To generate directed SF networks, we used the Barabási-Albert model [10], with the source and target of each node assigned randomly with equal probability.

A network before going through degeneration is referred here to as a parent network, whereas the degenerating descendants are denoted as their children networks. Each parent network consisted of 1, 000 neurons. To match the ratio of excitation and inhibition in the neocortex, 800 of them were randomly chosen to be excitatory neurons, whereas the remaining 200 neurons were chosen as inhibitory neurons. For each of the classes of networks we generated, we started with a connection probability of 10 %. The connection density and the size of the network clearly decreased as networks went through synaptic pruning and neurodegeneration. The parent networks were progressively degenerated and 10 stages of degeneration were selected for simulation. This was repeated 10 times to have variants of the parent networks and their degenerate children networks.

### Neuron model and simulation parameters

To simulate the spiking activity in the networks, each node was modelled as a leaky integrate-and-fire neuron and the neurons were interconnected using the current-based synapse model. The temporal evolution of the membrane potential of neuron *i*, denoted by *V*_*i*_, was governed by the following differential equation

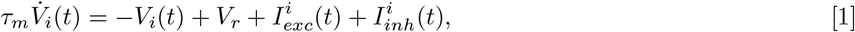

where *τ*_*m*_ is the membrane time constant, *V*_*r*_ is the resting membrane potential, and 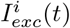 and 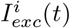 are the total excitatory and inhibitory synaptic inputs, respectively. When the membrane potential exceeded the spike threshold *θ*, the neuron elicited an action potential and the membrane potential was reset to *V*_*r*_ for 2 ms to mimic the refractoriness of biological neurons. The total excitatory or inhibitory current 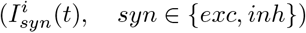 received by a neuron was given by:

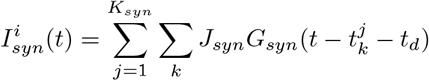

where *K*_*syn*_ is the number of inputs, 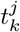 is the time of *k*^*th*^ spike from the *j*^*th*^ neuron, and *t*_*d*_ is the synaptic delay. Each neuron received *K*_*exc*_ and *K*_*inh*_ inputs from within the network and *K*_*ext*_ external excitatory inputs. The outer sum goes over the number of neurons and the inner sum goes over the spikes of each pre-synaptic neuron. *G*_*syn*_(*t*) is the synaptic kernel given by:

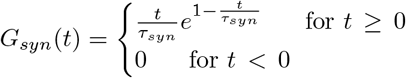

where *τ*_*syn*_ is the synaptic time constant, and *J*_*syn*_ is the amplitude of the synaptic kernel. *J*_*syn*_ is negative if the presynaptic neuron is inhibitory (*J*_*inh*_) and positive if it is excitatory (*J*_*exc*_). *t*_*d*_ is the synaptic delay. The ratio of total inhibitory and excitatory inputs was defined as 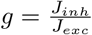 following previous results [11, 12].

To drive the neurons above their spike threshold, we stimulated each neuron with an external excitatory Poisson spike train with a rate of 6, 000 spikes/sec, corresponding to *K*_*ext*_ = 1000 inputs spiking each at 6 spikes/sec. To obtain different dynamical states of the network activity, two different values of relative inhibitory synaptic strength, *g* = 5 and *g* = 7, were used, which for an intact network corresponded to inhibition-dominant regimes. Neuron, synapse and network parameters are summarized in Table 1. All simulations were performed using the NEST simulator [42] and data was analysed using the Python programming environment (Version: 2.7).

**Table 1.** Neuron and network parameters.

**a. Neuron parameters**
| Symbol | Description | Value |
| --- | --- | --- |
| $\tau_m$ | Membrane time constant | 10 ms |
| $\theta$ | Threshold potential | -55 mV |
| $E_L$ | Equilibrium potential | -70 mV |
| $V_r$ | Reset potential | -70 mV |
| $t_{ref}$ | Refractory time | 2.0 ms |
| $t_d$ | Synaptic delay | 2.0 ms |
| $\tau_s$ | Synaptic time constant | 2.0 ms |

**b. Network parameters**
| Symbol | Description | Value |
| --- | --- | --- |
| $N_i$ | Network sizes | $\{i \times 100\}_{i=1}^{10}$ |
| $\rho_i$ | Network densities | $\{i\%\}_{i=0}^{10}$ |
| $N_{exc}$ | Fraction of exc. neurons | 80% |
| $N_{inh}$ | Fraction of inh. neurons | 20% |
| $prand$ | randomizing probability for SW network | 0.02 |
| $J_{exc}$ | excitatory weight | 0.1 mV |
| $g_j$ | inhibition to excitation ratio | $\{2.8, 4.8, 6.8\}$ |
| $J_{inh,j}$ | Inhibitory weights | $-g_j J_{exc}$ |

**c. Other parameters**
| Symbol | Description | Value |
| --- | --- | --- |
| $J_x$ | Poisson input weight | 0.1 |
| $p_{rate}$ | Poisson rate | 6 KHz |
| $simtime$ | Simulation time | 10s |

### Network degenerative models

#### Synaptic Pruning Strategies

We used five distinct synaptic pruning strategies; namely, *random, in-, out-, ordered* and *resilient* pruning strategies [13].

##### Random pruning

In this pruning strategy, each edge was randomly selected for deletion, unless it caused fragmentation. This was performed repeatedly until a last random spanning directed tree remained, where each edge was critical for keeping the network connected.

##### In-pruning

In this pruning strategy, we started by randomly selecting a node and systematically removed its incoming projections. Once the incoming edges of the chosen node were exhausted without fragmenting the network, another node was picked randomly to repeat the pruning procedure. Similar to random pruning, the pruning procedure was performed until a spanning tree was reached.

##### Out-pruning

In this pruning strategy, we started by randomly selecting a node and systematically removed its outgoing projections. Once the outgoing edges of the chosen node were exhausted without fragmenting the network, another node was picked randomly to repeat the pruning procedure. Similar to random pruning, the pruning procedure was performed until a spanning tree was reached.

##### Ordered maximum matching (ordered) pruning

This pruning strategy requires the knowledge of the full network connectivity. We first ordered the edges *E* of the network after exhaustive extraction of maximum matching sets. The first maximum matching (MM) set takes all edges into account, while the subsequent ones are maximum matching sets of the remaining edges, excluding the already extracted sets.

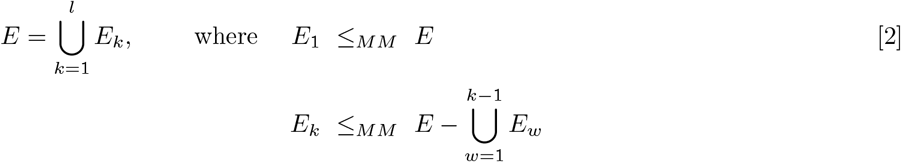

Hence, |*E*_*i*_| ≥ |*E*_*j*_| if *i* < *j*. Here, *A* ≤_*MM*_ *B* denotes “A is a maximum matching set of B”.

*E*_1_, …, *E*_*l*_ clearly form a partition of the set of all edges in the network. Each edge in the network belongs to some block, *E*_*k*_, in the partition. The relative rank, *rel. rank*, of an edge is then defined as the cardinality of the block it belongs to.

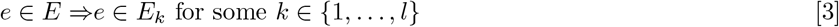

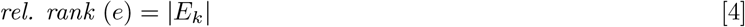

Deletion was performed in decreasing order of relative rank, i.e. all edges belonging to *E*_1_ were the first to be pruned in any order as long as they did not fragment the network. Then the procedure was repeated for all partitions according to the order of their indices.

*Resilient pruning*: This pruning strategy was aimed at keeping the controllability profile of the network [43] resilient to edge deletion. To this end, we designed three different methods. The first one was similar to the ordered-MM pruning, but in the reverse order, i.e. in a partition {*E*_1_, *E*_2_, …, *E*_*l*_} of *E*, where |*E*_*i*_| ≥ |*E*_*j*_| whenever *i* < *j*, the order of deletion was performed from *E*_*l*_ to *E*_1_. That is, the edges that were part of the maximum matching set corresponding to the original network were pruned only after all other edges were systematically removed. Thus, by definition, there was no change in the controllability profile of the network almost for the entire deletion process.

Both ordered pruning and resilient pruning required ordering all edges based on their relative rank (cf. eq. 3). According to the partitioning of edges as described in eq. 2, the cardinality of a maximum matching set determined the relative rank of the included edges, i.e. the higher the cardinality, the higher the rank order of the contained edges. While ordered-MM pruning was performed in decreasing order of relative rank, resilient pruning was performed in increasing order. Random pruning and out-pruning, however, required no knowledge of the relative rank of the network edges.

For each of the pruning strategies, we progressively pruned 10% of the existing synapses of networks at each step, until they eventually remained with just 1% connection density. At each stage, we had three different synaptic weight distributions, *g* ∈ {5, 7}, to represent the inhibitory-dominant regimes.

#### Neurodegenerative Strategies

The degree distribution defines the structure of the network best when all edges are of equal importance. Different degrees of nodes imply different levels of significance in the network. Spectral radius, centrality, enrichment, vertex cover, for instance, are all ultimately reflections of the degrees of the network nodes. Therefore, we devised one random and four degree-based neurodegenerative strategies. We refer to them as *random, increasing-out, increasing-degree, decreasing-degree* and *decreasing-out* degenerative strategies. Random neurodegeneration progressively deletes randomly selected neurons. Increasing-deg and decreasing-deg identify neurons with higher and lower degree, respectively and progressively deletes them in the respective order. Likewise, increasing-out and decreasing-out degenerative strategies were performed based on the out-degrees of the neurons.

In each neurodegenerative strategy, 100 neurons were removed at a time and the degenerative process was repeated until the last 100 neurons remained. In each removal of 100 neurons, the 4:1 proportion of excitatory and inhibitory population was maintained. That is, 80 excitatory and 20 inhibitory neurons were simultaneously removed at each stage of our neurodegenerative strategy. Similar to the synaptic pruning setting, each network was simulated for two different values of *g*, 5 and 7.

With 10 realizations, 3 network types, 2 relative inhibition scenarios, 2 degenerative schemes, 5 pruning or neurodegenerative strategies, and 10 stages of pruning, we generated a total of 60 parent networks and 5400 degenerates that made up a total of 5,460 networks. To discover the relationship between network structure and dynamics without the risk of bias on our degenerates, we simulated additional 3,600 networks with 4 realizations, 3 network types (ER, SW and SF),10 connection probabilities (1%-10%), 10 network sizes (100-1000), and 3 relative inhibitory synaptic strength *g* values 5, 6 and 7. Thus over 9,000 networks were included in this study.

### Measures of network activity dynamics

#### Firing rate (λ)

The mean firing rate of a neuron is the average number of action potentials it discharged per second (*λ*_*k*_). The firing rate of the network is then the mean firing rate of all neurons in the network. The variance of the firing rate is denoted by *σ*_*λ*_.

#### Coefficient of variation of inter-spike intervals (CV_ISI_)

To measure the irregularity of a spike train, we estimated the *CV* (*ISI*_*i*_) as:

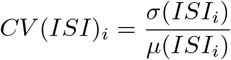

where *µ* and *σ* are the mean and the standard deviation of the inter-spike intervals of a neuron (*CV* (*ISI*_*i*_)). The mean of all these individual *CV* s is then used to estimate the level of spiking regularity in the network.

#### Pair-wise correlation (c)

The network spiking data was binned in 100 ms contiguous time windows to form vectors of spike counts for each neuron. A long spike train of neuron *i* can be binned into spike counts to form a vector of the form *n*_*i*_ = (*n*_*i*,1_, *n*_*i*,2_, …, *n*_*i*,_ *ℓ*), where *ℓ* is the number of bins. The pairwise correlation coefficient *c*_*i,k*_ of the firing patterns of neurons *i* and *k* is then given by:

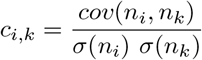

where *cov* and *σ* denote covariance and standard deviation. The level of correlation of a spiking network, denoted as 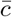 is estimated by the mean of all pairwise correlations coefficients in the network.

#### Fano-factor (FF)

To estimate the level of spike synchrony (including pairwise and all higher order correlations), we calculated the Fano factor of the network activity. The FF of the population activity is given by:

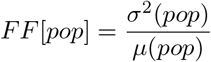

where *σ*^2^(*pop*) and *µ*(*pop*) denote the variance and mean of the population firing rate. To estimate the population firing rate vector (*pop*), we binned the spikes of all neurons in contiguous time bins (bin width = 100 ms). A population of independent Poisson processes yields a *FF* [*pop*] of unity and any mutual dependence among neurons would result in an increase of *σ*^2^[*pop*] and, hence, of *FF* [*pop*].

### Measures of network structure

#### Effective synaptic weight (ESW)

For a neuron *n*_*k*_ in a neuronal network with *nE*_*k*_ and *nI*_*k*_ numbers of presynaptic excitatory and inhibitory inputs, we define its effective synaptic weight as the net synaptic weight:

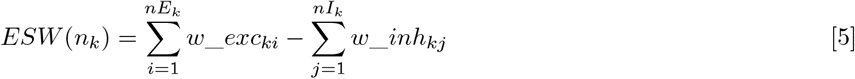

where *w*_*inh*_*kp*_ and *w*_*exc*_*kp*_ denote the connection weight from neuron *n*_*p*_ to neuron *n*_*k*_, depending on whether the input neuron is inhibitory or excitatory, respectively. Throughout our simulations, all excitatory and all inhibitory connections in a network had the same strength and obeyed the rule *J*_*inh*_ = − *gJ*_*exc*_. The average effective synaptic weight to neurons in the network was:

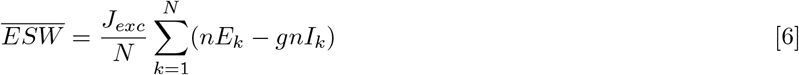

#### Pair-wise shared presynapses (shared)

The number of pair-wise shared presynapses in the network is the number of all pairs of neurons that have common presynaptic neighbours. For two neurons *n*_*i*_ and *n*_*j*_,*shared* (*n*_*i*_, *n*_*j*_), is the number of common presynaptic neurons.

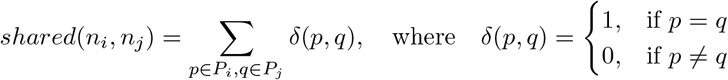

Here, *P*_*i*_ represents the set of all presynaptic neurons of neuron *n*_*i*_. The level of shared presynapses in the network, denoted by 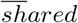, is the mean of all pairwise shared inputs.

### Measures of topology

#### Measure of randomness

To quantify the randomness of a network with a given connection density (*ρ*), we used the chi-squared test on its degree distribution *O*_*deg* with respect to the expected binomial degree distribution *E*_*deg* of the same connection density. We chose a critical chi-square value associated with *p* = 0.05 to accept or reject the null hypothesis. The number of degrees of freedom is one less than the size of the network.

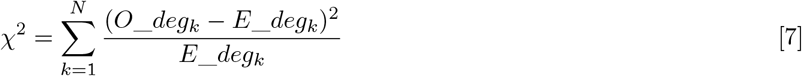

#### Measure of small-worldness

A number of measures have been suggested to estimate small-worldness of a network [44–47]. Here, we chose the *small-world propensity (ϕ)* [46] to check whether a network has strong small-world properties: short average path-length and high clustering coefficient because of its better classifying power. We computed *ϕ* for all our networks at different stages of pruning to measure how pruning changed the small-world property.

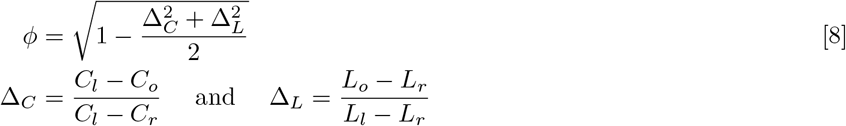

Here, *L* and *C* denote the average characteristic path length and the average clustering coefficient of the network under consideration. While the subscripts *o* denotes the candidate network, *l* and *r* represent the ring lattice and random networks of equal density as the candidate network. For a network to exhibit strong small-world properties, its *ϕ* should at least be 0.6 [46].

#### Measure of scale-freeness

To check to what extent a network is scale-free, we fitted its degree distribution to a power law (*p*(*k*) = *ck*^−*γ*^). The fitting exponent *γ* determines whether the network is scale-free or not. If *γ* is bounded between 2 and 3, it exhibits scale-free features [10].

## ACKNOWLEDGMENTS

This work was funded in part by: the EU Erasmus Mundus Joint Doctorate Program EUROSPIN, the Carl-Zeiss Foundation, the Swedish Research Council (Research Project Grant, StratNeuro, India-Sweden collaboration grants to Arvind Kumar). We thank Prof. Stefan Rotter and Dr. Michelle Rudolph-Lilith for helpful discussions.

## Supporting information

**S1.**
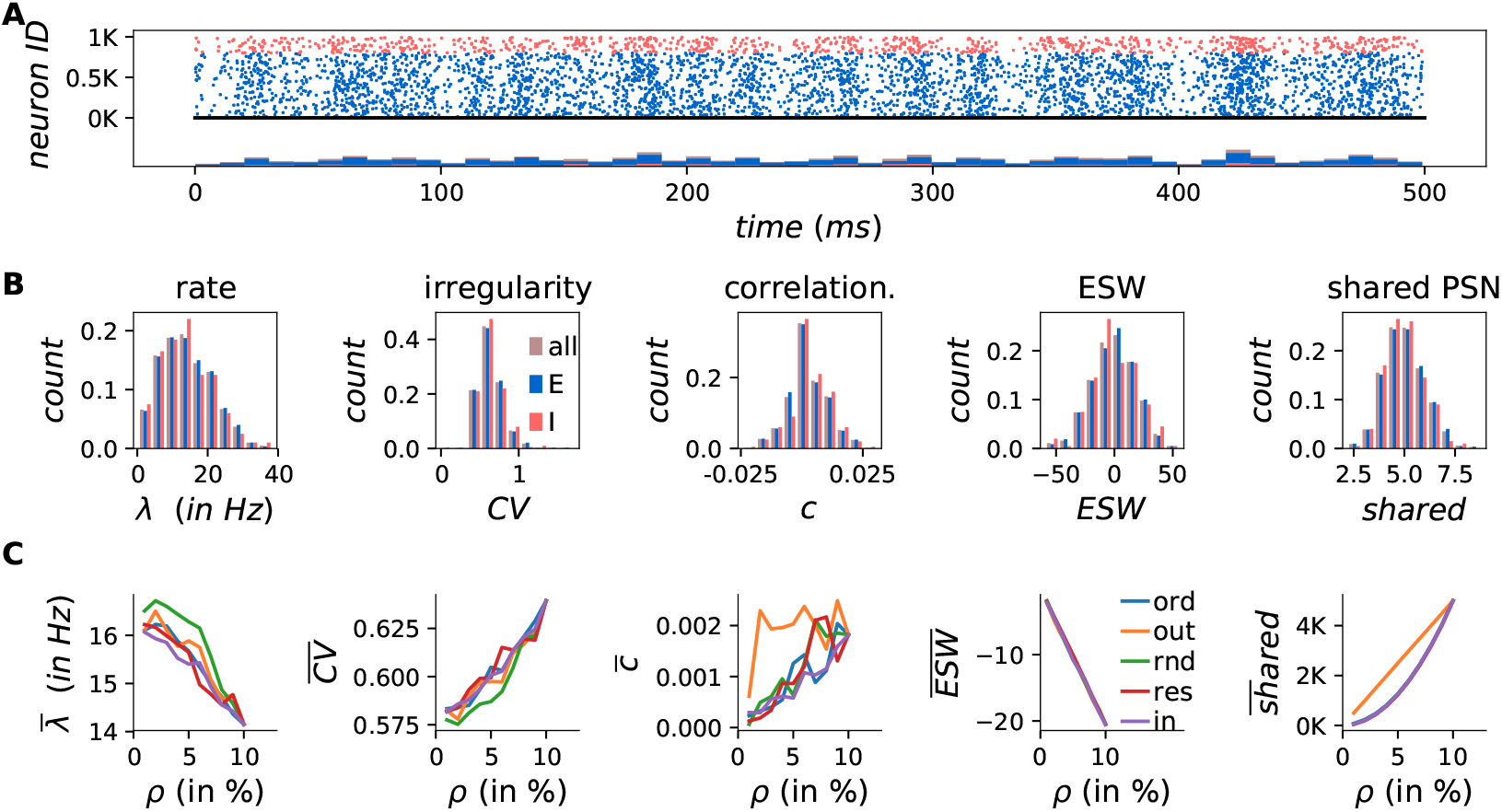
Spiking activity statistics of an Erdös-Rényii network under synaptic pruning. The ratio of the number of excitatory and inhibitory neurons was 4. Synapses from inhibitory neurons were 5 times stronger than those from excitatory neurons (g=5). A. Spiking activity for 1s and the corresponding Peri-Stimulus Time Histogram (PSTH) of the network of 1,000 neurons. B-D. Sample dynamical quantities for an unpruned ER spiking network: firing rate (A), spiking irregularity via coefficient of variation of the inter-spike intervals (B) and mean pairwise correlation of neurons. E-F. Sample structural quantities: effective synaptic weight (E) and the percentage of shared presynaptic neighbours of neurons (F). G-K. The mean of each of the quantities in (B-F) as the network loses its 10K synapses at a time until the last 10K synapses remained. The five pruning strategies are ordered (blue), out-(yellow), random (green), resilient (red) and in-(purple) pruning strategies. Since the pruning graphs are shown in increasing connection density (*ρ*), pruning progressed from right to left.

**S2.**
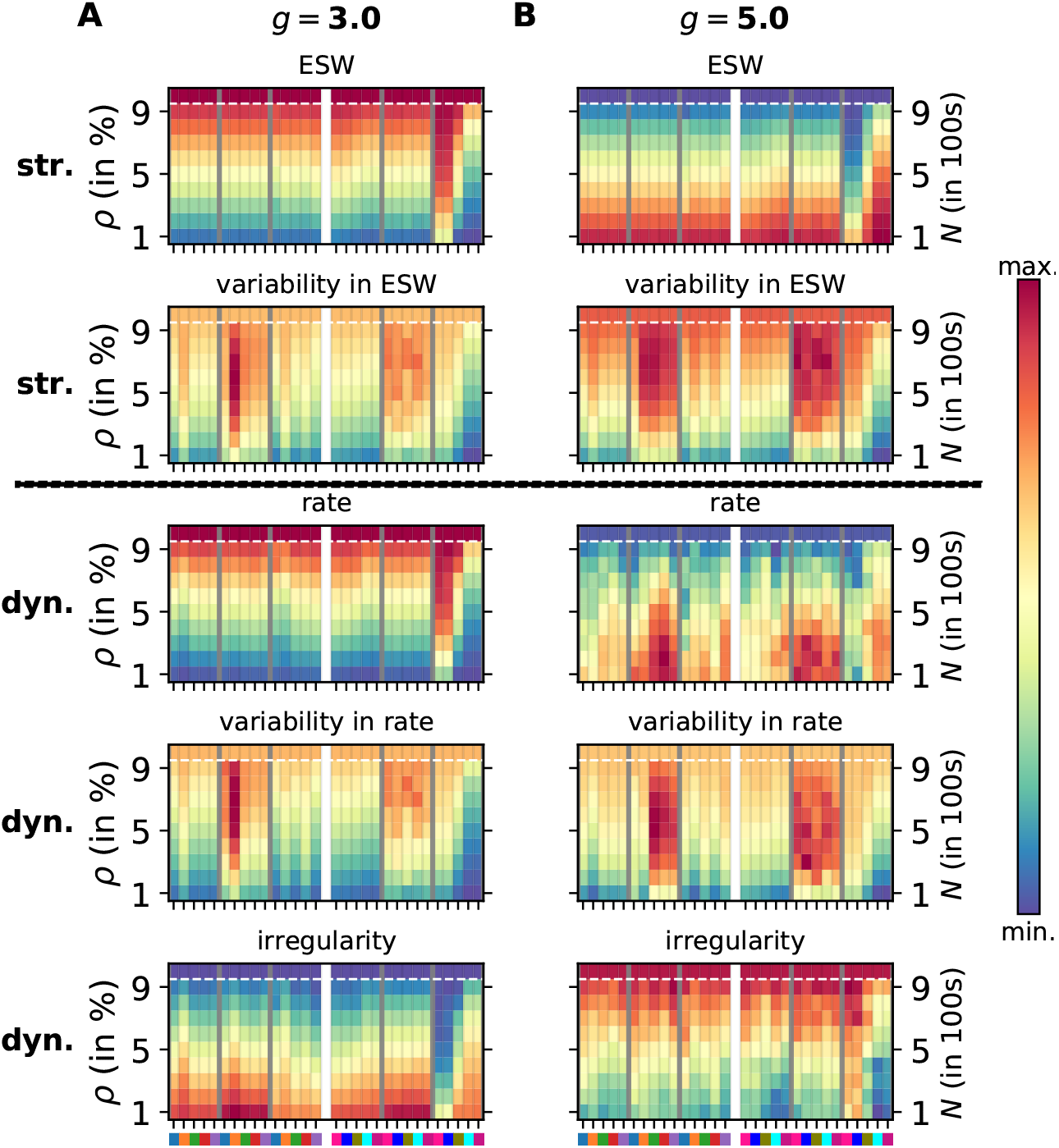
Comparison of structural and dynamical features of spiking networks with progressing degeneration. Each panel is presented in the same way as in Fig. 1. The top two rows comprise structural measures: effective synaptic weight (*ESW*) and its variability, the bottom three rows represent dynamical measures: firing rate (*λ*), its variability (*σ*_*λ*_), and the coefficient of variation of inter-spike intervals (*CV*). A. Excitation-dominant state (*g* = 3.) B. Inhibition-dominant state (*g* = 5.)

